# ARTEMIS - a method for topology-independent superposition of RNA 3D structures and structure-based sequence alignment

**DOI:** 10.1101/2024.04.06.588371

**Authors:** Davyd R. Bohdan, Janusz M. Bujnicki, Eugene F. Baulin

**Author notes:** corresponding author, correspondence may be also sent to: Janusz M. Bujnicki, Eugene F. Baulin.

## Abstract

Non-coding RNAs play a major role in diverse processes in living cells with their sequence and spatial structure serving as the principal determinants of their function. Superposition of RNA 3D structures is the most accurate method for comparative analysis of RNA molecules and for inferring sequence alignments. Topology-independent superposition is particularly relevant, as evidenced by structurally similar RNAs with sequence permutations such as tRNA and Y RNA. To date, state-of-the-art methods for RNA 3D structure superposition rely on intricate heuristics, and the potential for topology-independent superposition has not been exhausted. Recently, we introduced the ARTEM method for unrestrained pairwise superposition of RNA 3D modules and now we developed it further to solve the global RNA 3D structure alignment problem. Our new tool ARTEMIS significantly outperforms state-of-the-art tools in both sequentially-ordered and topology-independent RNA 3D structure superposition. Using ARTEMIS we discovered a helical packing motif to be preserved within different backbone topology contexts across various non-coding RNAs, including multiple ribozymes and riboswitches. We anticipate that ARTEMIS will be essential for elucidating the landscape of RNA 3D folds and motifs featuring sequence permutations that thus far remained unexplored due to limitations in previous computational approaches.

## INTRODUCTION

Non-coding RNAs play a major role in diverse processes in living cells with their sequence and spatial structure serving as the principal determinants of their function [1]. The rigid-body superposition of RNA 3D structures stands as a fundamental method for the comparative analysis of RNA molecules [2]. By superimposing structures of homologous RNAs, it becomes possible to discern conserved and variable segments [3], which is commonly used in template-based RNA structure modeling [4]. The 3D superposition of distinct structures of RNA molecules proves beneficial in studies of functionally-related dynamics, such as those between different stages of self-splicing of introns [5] or between apo- and holo- states of riboswitches [6]. Furthermore, RNA 3D structure superposition is invaluable for inferring correct sequence alignments [7], given that RNA exhibits significantly less conservation at the sequence level compared to its structural conservation [8]. Although flexible superposition is superior for deriving sequence alignments [9], it poses a more challenging task; hence, rigid-body 3D structure alignment remains the gold standard approach [2].

Besides comparisons centered on sequentially-ordered RNA molecules, comparisons have to take into account molecules that are related but don’t keep the same structure topology and exhibit sequence permutations: Several viral and bacterial RNAs have been identified to mimic tRNA structures, exhibiting both circularly and non-circularly permuted sequence matchings [10, 11]. Circular permutations have been observed in various natural non-coding RNAs [12, 13]. Recently, a natural non-circularly permuted variant of a non-coding RNA, specifically the hammerhead ribozyme, was reported [14]. Moreover, topology-independent superposition proves valuable for analyzing G-quadruplexes, which adopt diverse backbone topologies in both RNA and DNA [15, 16], given the rapidly increasing number of discoveries revealing their relevance in human diseases and therapeutics [17, 18].

While several methods, such as Rclick [7] and a more recent development RNAhugs [19], demonstrate capability in topology-independent superposition, comprehensive benchmarking of their performance in this regard has not been undertaken, to the best of our knowledge. Moreover, US-align, standing as the state-of-the-art tool for sequentially-ordered superposition, also exhibits proficiency in topology-independent superposition, although this aspect was not addressed in the original paper [2]. The greatest challenge of the RNA 3D structure alignment problem, both sequentially-ordered and topology-independent, is the exponential time complexity of the computational problem [2]. Currently, this is being handled either by employing fast but simple heuristics [2] or by relying on intricate but slow ones [19]. Furthermore, the problem of detecting backbone-permuted RNA 3D structure similarities has not been previously explored.

Recently, we developed the ARTEM method for unrestrained pairwise superposition of arbitrary RNA 3D modules [20]. In this work, we present ARTEMIS, an application of ARTEM methodology to address the global RNA 3D structure alignment problem. The heuristic behind ARTEMIS operates in polynomial time and ensures the optimal solution, provided it includes at least one residue-residue match with a near-zero RMSD, a condition commonly met in RNA structures due to their characteristic recurrent interactions [20]. ARTEMIS significantly outperforms state-of-the-art tools in both sequentially-ordered and topology-independent RNA 3D structure superposition. Leveraging ARTEMIS, we discovered a helical packing motif to be preserved in different backbone topology contexts in diverse non-coding RNAs, including multiple ribozymes and riboswitches.

## MATERIALS AND METHODS

### RNA 3D structure alignment problem

We define the computational problem of pairwise RNA 3D structure alignment as follows. Let *X* and *Y* denote two RNA structures of *N* and *M* residues, respectively. Without loss of generality, we assume *N ≥ M* and consider *X* as the static *reference structure,* with *Y* serving as the *query structure* undergoing superposition on *X*. An alignment (or superposition) between *X* and *Y* is then defined as a subset of residue pairs *(X_i_, Y_j_)*, with the condition that each residue can only have one match at most. If an alignment includes at least two pairs of residues *(X_i_, Y_j_)* and *(X_k_, Y_l_)* such that *i < k* and *j > l*, it is termed a *topology-independent alignment*. In other words, it represents a superposition where residue-residue matchings cannot be sorted monotonically for both sequences simultaneously. Conversely, if such a condition is not met, the superposition denotes *a sequentially-ordered alignment*, or simply a sequence alignment in the traditional sense.

Among all possible alignments, our focus lies on identifying the alignment that captures the most structurally similar subset of residues between *X* and *Y* while achieving the highest possible *coverage*, defined as the ratio of aligned residues to sequence length. In this context, structural similarity refers to the resemblance between rigid coordinate models, commonly assessed through the *root mean square deviation* (*RMSD*) [21]. *TM-Score* metric [22] for proteins and its variant *TM-Score_RNA_* [23] for RNAs enable the consideration of both coverage and structural similarity.

*TM-Score_RNA_* positively depends on the *L_ali_* length of the alignment (the number of matched residue pairs), inversely depends on the *d_i_* distances between the *C3′* atoms of the matched residues, and is normalized by the length *L* of one of the chains, ensuring that the score falls within the range of zero to one [2]. For each superposition, two *TM-Score_RNA_* values can be computed: *TM1-Score_RNA_*, normalized by the longer chain length *N*, and *TM2-Score_RNA_*, normalized by the shorter chain length *M*. Likewise, two coverage values are provided for each superposition: *L_ali_ / N* and *L_ali_ / M*.

We define the pairwise RNA 3D structure alignment problem as the task of identifying the optimal alignment, whether sequentially-ordered or with potential sequence permutations (topology-independent), which maximizes the sum of (*TM1-Score_RNA_* + *TM2-Score_RNA_*) values.

Drawing from the well-established field of protein bioinformatics, where a 3D fold is defined as the spatial arrangement of all secondary structure elements without considering the sequential order or the connecting loops, in this work we employ a similar conceptual framework for RNA molecules. Here, the RNA 3D fold emphasizes the spatial arrangement of sequence segments in RNA, involved in base-paired structured motifs, primarily helices, while disregarding the order in which these paired sequence segments appear or the nature of the intervening sequences that link them. A common 3D fold implies geometric similarity of RNA structures, without implying the same topology of the chain. Molecules exhibiting the same 3D fold may be related evolutionarily (homologs) or may arise through convergent evolution (analogs).

### ARTEMIS algorithm

To identify the optimal alignment, we developed the ARTEMIS algorithm (using **ARTEM** [20] to **I**nfer **S**equence alignment), which operates under the assumption that the ideal superposition involves at least one pair of matched residues exhibiting a near-zero RMSD.

ARTEMIS superimposes the query structure *Y* on the reference *X* across all possible residue pairs between the structures (Supplementary Figure S1, lines 20-32). Each superposition is computed using the Kabsch algorithm [21], employing a 3-atom representation of the residues. This representation comprises three pseudo atoms corresponding to the position of the phosphate group (center of mass of *OP1* and *O5′* atoms), the ribose (center of mass of *C2′* and *C4′* atoms), and the base (glycosidic base atom). The selection of these pseudo atoms was determined during preliminary experiments. For each superposition, ARTEMIS identifies the *hit*, which represents the set of pairs of mutually closest residues between the structures, with the distance between their *C3′* atoms falling below the *MATCHRANGE1* threshold of *3.5 Å* (Supplementary Figure S1, lines 26-29).

For the *TOPLARGEST = M* largest hits, ARTEMIS constructs extended matchings (Supplementary Figure S1, lines 33-53). For each hit, the query structure *Y* undergoes re-superimposition on *X* using the 3-atom representations of the hit’s matched residues. Subsequently, the adjusted superposition is utilized to identify an extended set of mutually closest residues between *X* and *Y*, employing the *MATCHRANGE2* threshold of *8 Å* (Supplementary Figure S1, lines 40-43). The resultant matching serves as a candidate topology-independent alignment.

The same adjusted superposition is subsequently utilized to determine the sequentially-ordered alignment (Supplementary Figure S1, lines 45-49). For this purpose, a scoring matrix for the Needleman-Wunsch global sequence alignment algorithm [24] is prepared using the matrix of pairwise C3′-C3′ distances (Figure 1A). This matrix is constructed by negating the distance values (Figure 1B) and then adding the largest distance between the matched residues, denoted as *SHIFT1*, to all values in the negative matrix (Figure 1C), along with an additional term *SHIFT2 = 3 Å* (Figure 1D). Subsequently, the Needleman-Wunsch algorithm derives the optimal sequence alignment to maximize the score. A higher *SHIFT2* parameter results in increased coverage, while a *SHIFT2* value of *0 Å* ensures that the sequentially-ordered alignment remains a subset of the corresponding topology-independent alignment.

**Figure 1.**
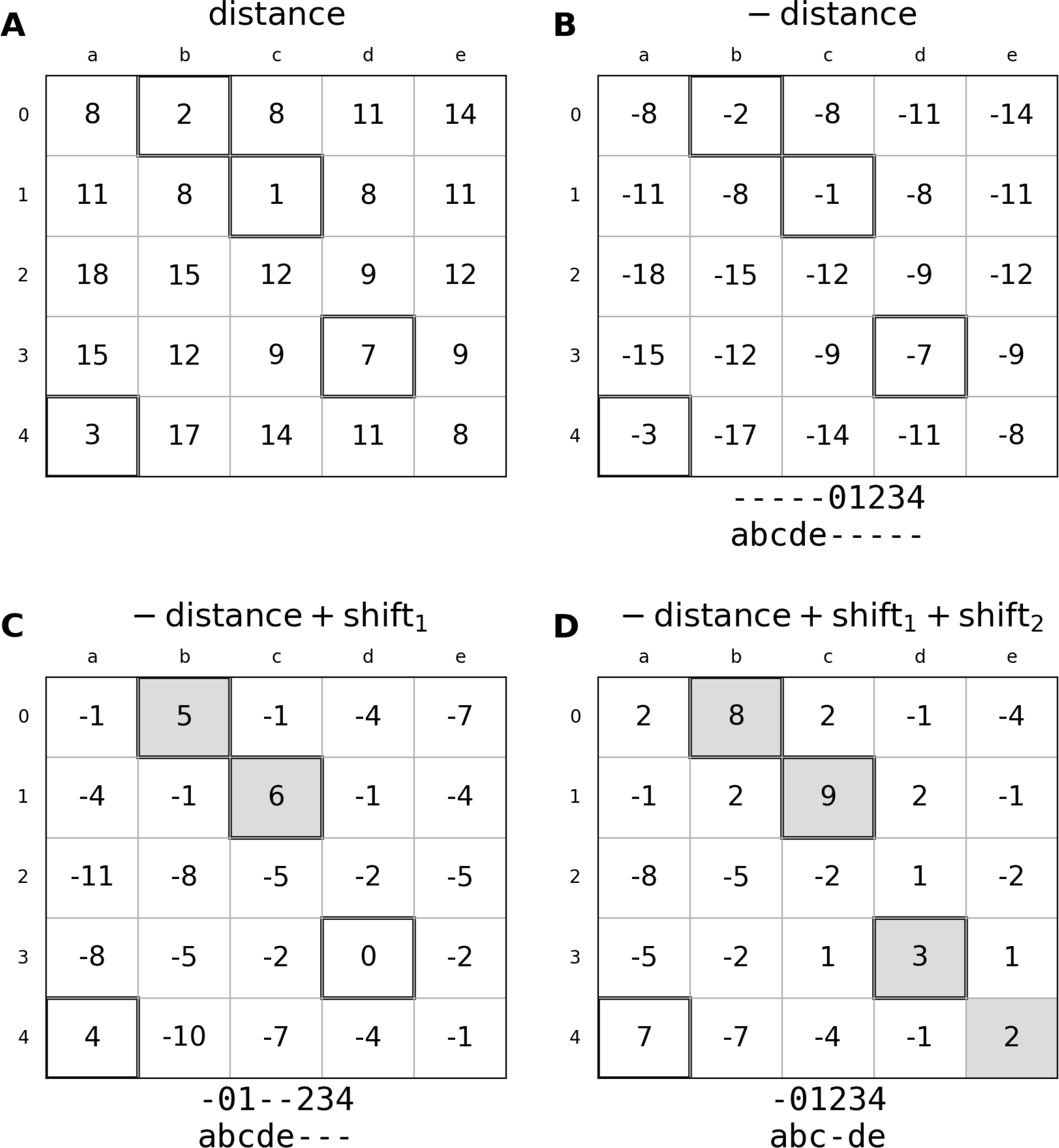
A demonstration example of the scoring matrix preparation procedure used in ARTEMIS for constructing sequentially-ordered alignments. (A) A matrix of C3′-C3′ distances. (B) The matrix of negative distances. (C) The matrix after adding the SHIFT1 term. (D) The matrix after adding the SHIFT2 term. Solid-framed cells denote the initial topology-independent hit, while gray-filled cells indicate sequentially-ordered alignments, visually represented as gapped lines below each matrix.

ARTEMIS ultimately provides the optimal sequentially-ordered alignment alongside the optimal topology-independent alignment that encompasses sequence permutations (Supplementary Figure S1, lines 54-55). It’s known [2] that, for a given alignment, the minimum-RMSD superposition obtained through the Kabsch algorithm does not typically maximize the TM-Score_RNA_ value. Consequently, for each candidate alignment, ARTEMIS iteratively splits L_ali_ matched residue pairs into fragments of lengths L_ali_ / 2, L_ali_ / 4, …, 3. It proceeds to superimpose each fragment utilizing the Kabsch algorithm, selecting residue pairs whose *C3′-C3′* distance value falls below *d0(M)*. Subsequently, the Kabsch superposition of all matched residue pairs is executed based on the selected pairs, and the value of *(TM1-Score_RNA_ + TM2-Score_RNA_)* is calculated. The *TM1-Score_RNA_* and *TM2-Score_RNA_* yielding the maximum sum are then assigned to the candidate alignment.

Similar to ARTEM [20], the theoretical time complexity of ARTEMIS is *O(N^2^ * M^2^),* given that ARTEMIS computes *N*M* pairwise distances to determine the set of mutually closest residues for each of the *N*M* single-residue seed superpositions (Supplementary Figure S1, lines 20-30).

### ARTEMIS algorithm acceleration

To expedite ARTEMIS when the length *M* of the smaller structure *Y* exceeds 500 nucleotides, a faster heuristic is employed. Initially, we constrain the total number of single-residue seed superpositions. Instead of considering all *N*M* possible pairs, ARTEMIS traverses only each *STEP = 1 + M // STEPDIV* residue of *X*, with *STEPDIV = 100* by default. Additionally, we adjust the *TOPLARGEST* value to twice the number of machine CPUs utilized by the ARTEMIS program. Furthermore, we elevate the *SHIFT2* value to *20 Å*. This adjustment does not impact speed but enhances alignment coverage by better capturing related but flexible regions between two long chains within the Needleman-Wunsch alignment. These modifications enable ARTEMIS to process large RNA chains efficiently while maintaining alignment quality.

### ARTEMIS tool implementation

ARTEMIS was implemented in Python3 as a command line interface (CLI) application and is available at https://github.com/david-bogdan-r/ARTEMIS. It requires two DNA- or RNA-containing files in PDB or mmCIF format as input, along with optional arguments, including *MATCHRANGE, TOPLARGEST, SHIFT2*, and *STEPDIV* as described above. The standard output of ARTEMIS comprises information on the input structures, pairwise sequence alignment, and essential alignment quality metrics such as TM-score_RNA_, RMSD, and alignment length L_ali_. Details of topology-independent alignment are included in the output either upon user request or if its TM2-Score_RNA_ surpasses that of the sequentially-ordered alignment by at least 10%. Optionally, users can save the coordinates of the superimposed query structure along with a list of matched residues and the distances between them for either alignment. ARTEMIS is designed as a parallelized program and utilizes all available processors on the machine by default.

ARTEMIS is available to be adapted as a Python3 module, facilitating its integration into environments such as Jupyter Notebooks (https://github.com/david-bogdan-r/ARTEMIS/blob/main/demo.ipynb). Independent implementations of the RNA structure class and the primary ARTEMIS procedure class enable users to bypass unnecessary preliminary procedures, particularly when aligning multiple query structures to a single reference.

### Benchmarking

The performance evaluation of ARTEMIS was based on a dataset comprising 637 RNA chains [2], previously employed for benchmarking US-align [2], RMalign [25], STAR3D [26], ARTS [27], and Rclick [7]. To compare sequentially-ordered alignments, we conducted superpositions by ARTEMIS and US-align and utilized the alignment quality metrics values of the other tools from the prior benchmark [2]. The results of US-align obtained by us and those from the previous benchmark were verified to be identical. Assessment of sequentially-ordered superposition quality was performed on 168,916 structure pairs, as for the remaining 33,650 pairs, at least one of the tools failed to provide a superposition [2].

For topology-independent alignment comparisons, we utilized standalone versions of US-align, Rclick, and RNAhugs (Supplementary Table S1). For RNAhugs, we considered separately the geometric (GEOS) and genetic (GENS) modes. Both GEOS and GENS inherently perform topology-independent superposition by default. Due to RNAhugs’s potentially lengthy runtime, which can be tens of seconds even for a superposition of two RNAs under 100 residues, we excluded it from the primary benchmark. As Rclick and RNAhugs do not provide *TM-Score_RNA_* values, we directed their superpositions to US-align to calculate the values without re-superimposing the structures (Supplementary Table S1). Subsequently, we considered RMSD, L_ali_, and TM-Score_RNA_ values as reported in US-align outputs. Evaluation of topology-independent superposition quality was conducted on all 202,566 structure pairs.

Execution time comparisons were conducted for 151,735 structure pairs for sequentially-ordered superposition and 184,282 structure pairs for topology-independent superposition. These pairwise superpositions were executed on an Intel Core i7-8750H machine equipped with 12 CPU cores and 16 GB RAM. For the remaining 17,181 and 18,284 structure pairs, respectively, ARTEMIS was run on a faster machine to speed up the process.

For each benchmark, we also included the “best competitor” column, determined by selecting, for each structure pair, the alignment with the highest combined *(TM1-Score_RNA_ + TM2-Score_RNA_)* value among all tools except ARTEMIS. To evaluate the statistical significance of performance differences among the tools, we employed the paired Student’s t-test, implemented in the SciPy Python library (https://docs.scipy.org/doc/scipy/reference/generated/scipy.stats.ttest_rel.html).

To identify similar folds exhibiting sequence permutations, we compared sequentially-ordered and topology-independent superpositions generated by ARTEMIS. We defined a pair of structures to exhibit a backbone-permuted similarity if the topology-independent TM1-Score_RNA_ value exceeded the sequentially-ordered TM1-Score_RNA_ value by at least 0.1.

## RESULTS

### Benchmarking ARTEMIS performance

We compared ARTEMIS with state-of-the-art tools in their performance on the curated dataset of 637 RNA chains previously utilized for benchmarking US-align. ARTEMIS significantly outperformed the other tools in terms of TM-*Score_RNA_* values, both in constructing sequentially-ordered alignments (Figure 2, Figure 3, Supplementary Table S2) and topology-independent alignments (Figure 4, Supplementary Table S3).

**Figure 2.**
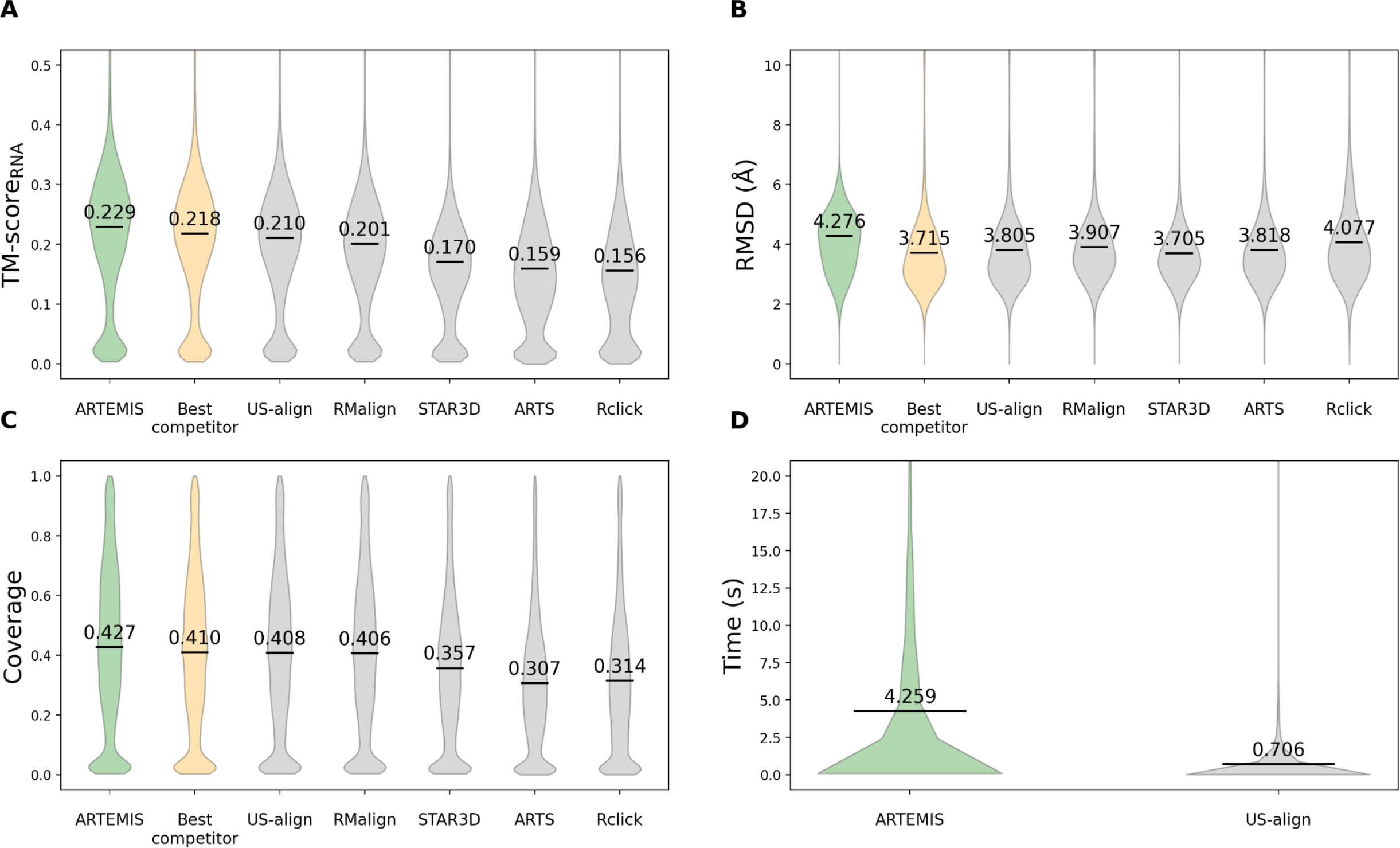
Sequentially-ordered alignment benchmark results. Performance metrics include (A) TM-score_RNA_, (B) RMSD, (C) coverage, and (D) execution time. ARTEMIS demonstrates superior performance in terms of TM-score_RNA_, while reporting slightly higher RMSD values, attributed to higher coverage values. Time measurements were conducted for ARTEMIS and US-align exclusively, as the results of other tools were directly obtained from the previous benchmark [2].

**Figure 3.**
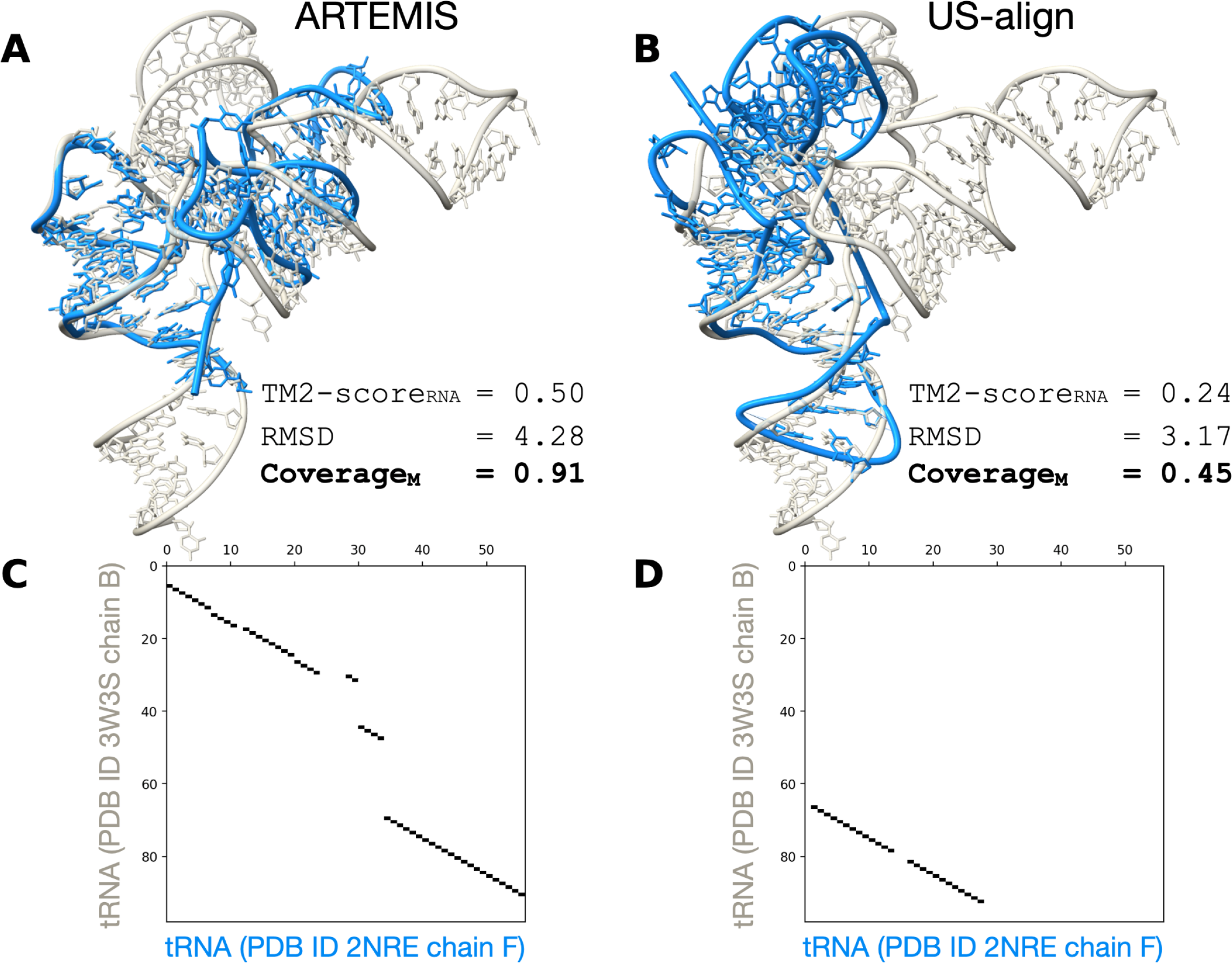
A demonstration example showcasing ARTEMIS’ superior performance (A, C) compared to US-align (B, D). Panels C and D depict alignment plots corresponding to the structure superpositions shown in panels A and B, respectively.

**Figure 4.**
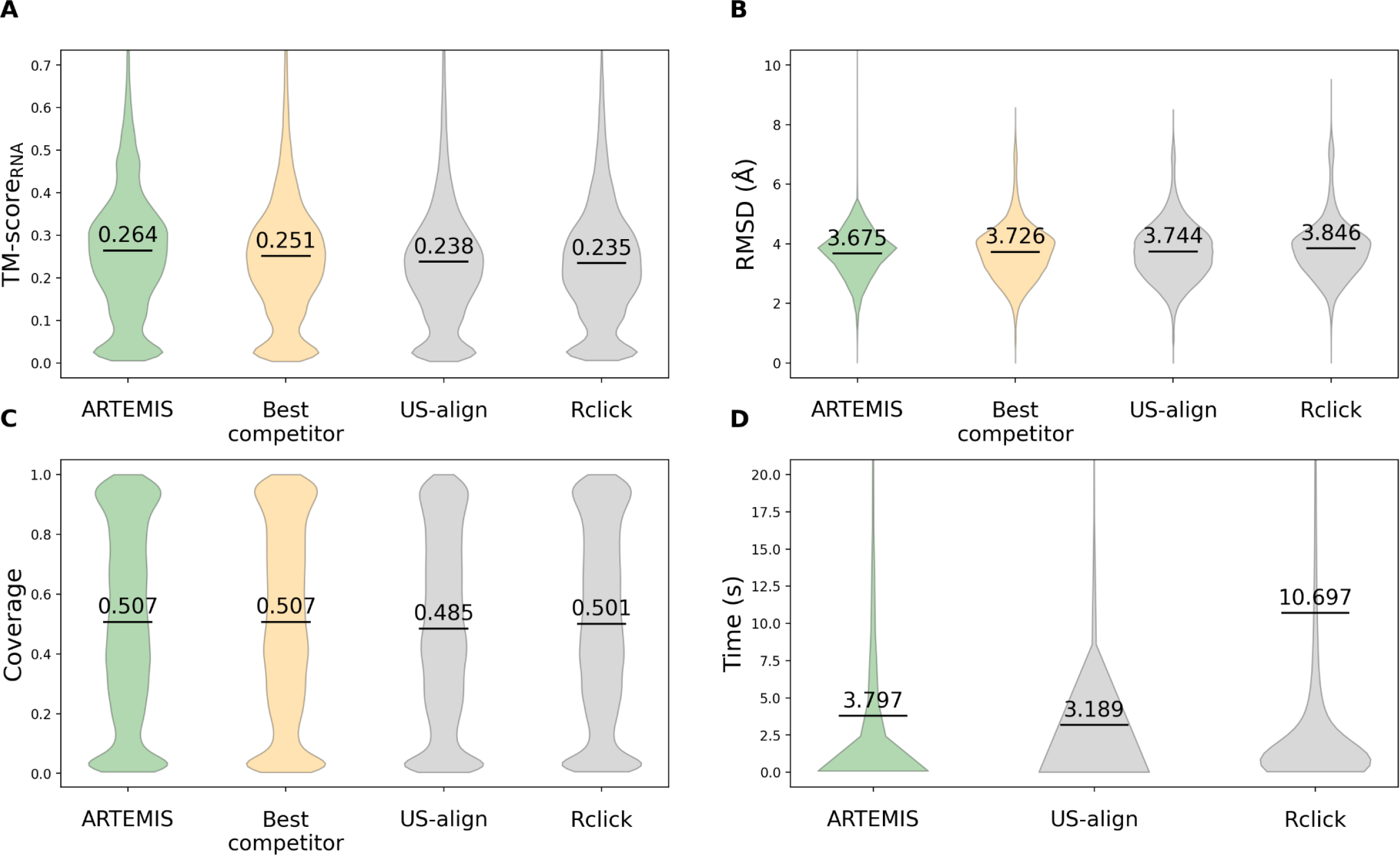
Topology-independent alignment benchmark results. Performance metrics include (A) TM-score_RNA_, (B) RMSD, (C) coverage, and (D) execution time. ARTEMIS demonstrates superior performance compared to existing tools in terms of TM-score_RNA_, with lower RMSD values and higher coverage values.

Compared to US-align, thus far the golden standard, ARTEMIS constructed sequentially-ordered alignments with superior TM-Score*_RNA_* values (mean 0.229 vs. mean 0.21, Figure 2A) and reported slightly higher RMSD values (mean 4.276 Å vs. mean 3.805 Å, Figure 2B) attributable to higher coverage values (mean 0.427 vs. mean 0.408, Figure 2C). Although US-align operates faster than ARTEMIS in the sequentially-ordered alignment mode (mean 0.706 s vs. mean 4.259 s, Figure 2D), ARTEMIS still required only four seconds on average, with the maximum time for a superposition of two ribosomal RNAs being under four minutes.

In topology-independent superposition, ARTEMIS (mean TM-Score*_RNA_*0.264, Figure 4A) surpasses US-align and Rclick (0.238 and 0.235, respectively). In this case, ARTEMIS also reports lower RMSD values (Figure 4B) and higher coverage values (Figure 4C). The execution times of ARTEMIS and US-align are comparable, whereas Rclick is three times slower (Figure 4D).

### Analysis of tRNA-like structures

Within the benchmark dataset, we identified 97 pairs of similar RNA 3D folds with sequence permutations (Supplementary Table S4). Among these pairs, 80 featured a bacterial Y RNA molecule (PDB entry 6CU1, chain A) [11] forming circularly permuted superpositions with various tRNAs (Figure 5). Subsequently, we manually collected a representative set of a tRNA molecule [28] and five tRNA-like structures (tmRNA fragment [29], selenocysteine tRNA [30], tRNA-like structure from Turnip Yellow Mosaic Virus [10], bacterial Y RNA [11], and tRNA-like structure from Brome Mosaic Virus [31]) and evaluated their pairwise topology-independent superpositions constructed by ARTEMIS, US-align, Rclick, and RNAhugs (Table 1, Supplementary Table S5).

**Figure 5.**
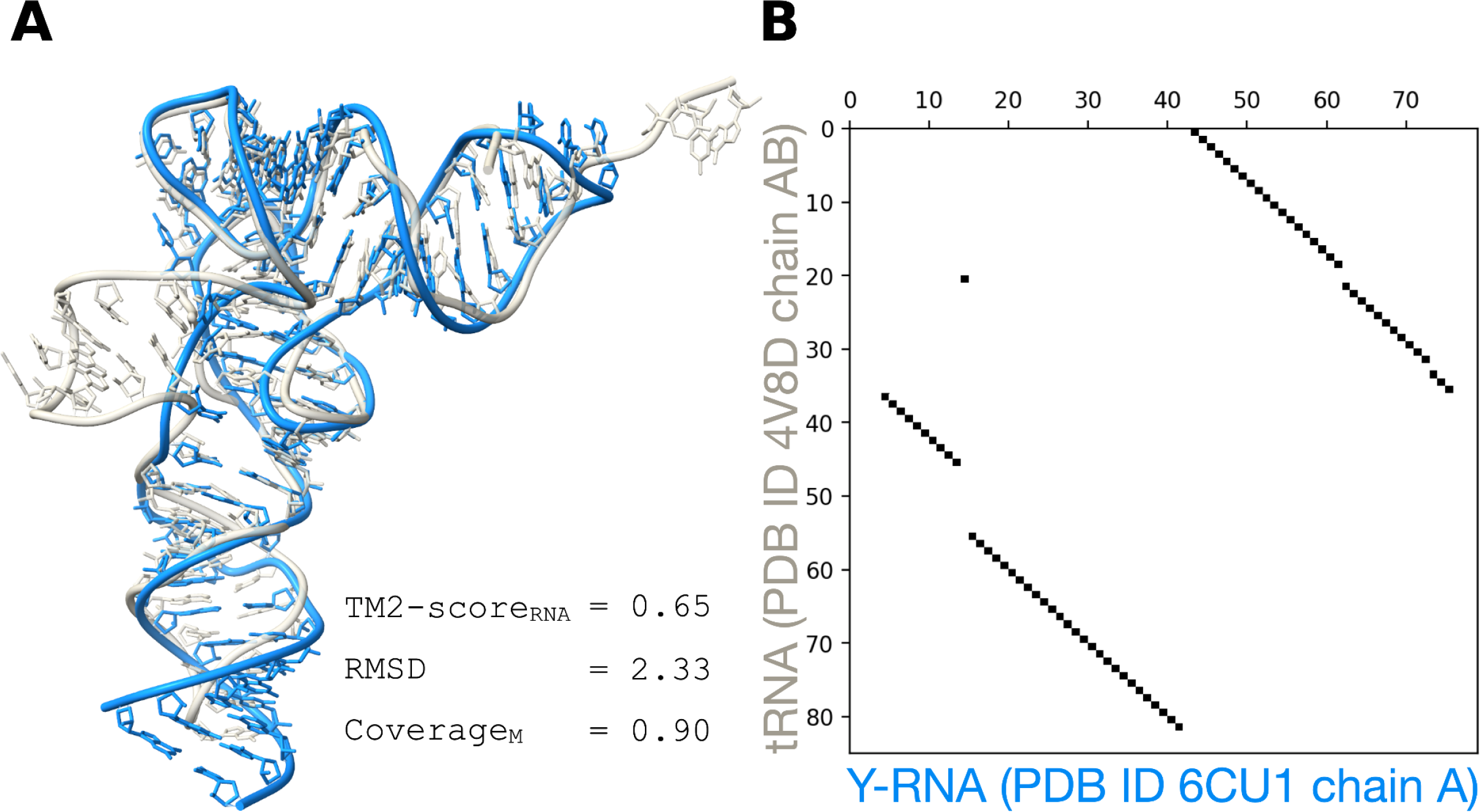
An illustration of a circularly permuted superposition between a bacterial Y RNA (PDB entry 6CU1, chain A) and a tRNA (PDB entry 4V8D, chain AB), identified by ARTEMIS. (A) 3D structure superposition. (B) Alignment plot.

**Table 1.** Performance of the topology-independent superposition tools on tRNA-like structures.

| Tool |  |  |  | ARTEMIS |  | US-align |  | Rclick |  | RNAhugs<br>GEOS |  | RNAhugs<br>GENS |  |
| --- | --- | --- | --- | --- | --- | --- | --- | --- | --- | --- | --- | --- | --- |
| tRNA vs. | PDB id<br>Chain | Length | Permutation | TM1<br>score | TM2<br>score | TM1<br>score | TM2<br>score | TM1<br>score | TM2<br>score | TM1<br>score | TM2<br>score | TM1<br>score | TM2<br>score |
|  | 1ivsC | 75 |  |  |  |  |  |  |  |  |  |  |  |
| tmRNA<br>fragment | 2czjB | 62 | no | 0.39 | 0.435 | 0.365 | 0.413 | 0.34 | 0.379 | 0.328 | 0.351 | 0.339 | 0.376 |
| tRNA-Sec | 3addC | 88 | no | 0.453 | 0.485 | 0.461 | 0.494 | 0.445 | 0.477 | 0.446 | 0.473 | 0.446 | 0.473 |
| TYMV<br>tRNA-like | 4p5jA | 83 | non-circular | 0.45 | 0.468 | 0.464 | 0.481 | 0.407 | 0.416 | 0.43 | 0.44 | 0.421 | 0.43 |
| Y RNA | 6cu1A | 79 | circular | 0.613 | 0.632 | 0.613 | 0.633 | 0.612 | 0.631 | 0.554 | 0.566 | 0.546 | 0.556 |
| BMV<br>tRNA-like | 7samA | 169 | non-circular | 0.261 | 0.372 | 0.21 | 0.3 | 0.273 | 0.401 | 0.244 | 0.378 | 0.196 | 0.291 |
| Mean |  |  |  | 0.433 | 0.478 | 0.423 | 0.464 | 0.415 | 0.461 | 0.4 | 0.442 | 0.39 | 0.425 |

ARTEMIS demonstrated superior *TM-score_RNA_* values, although all tools performed very closely (Table 1). Interestingly, all tools generated almost identical superpositions for all structures except the tRNA-like structure from Brome Mosaic Virus (BMV, PDB entry 7SAM, chain A). The BMV structure includes two tRNA-mimicking fragments, with helical fragment A mimicking the tRNA acceptor stem and helical fragment B3 mimicking the tRNA anticodon stem [31]. These two fragments cannot be superimposed on a tRNA structure simultaneously due to their extended arrangement. ARTEMIS, Rclick, and RNAhugs(GEOS) correctly superimposed fragment A onto the acceptor stem, while US-align correctly superimposed fragment B3 onto the anticodon stem. RNAhugs(GENS) reported an incorrect superposition (Figure 6).

**Figure 6.**
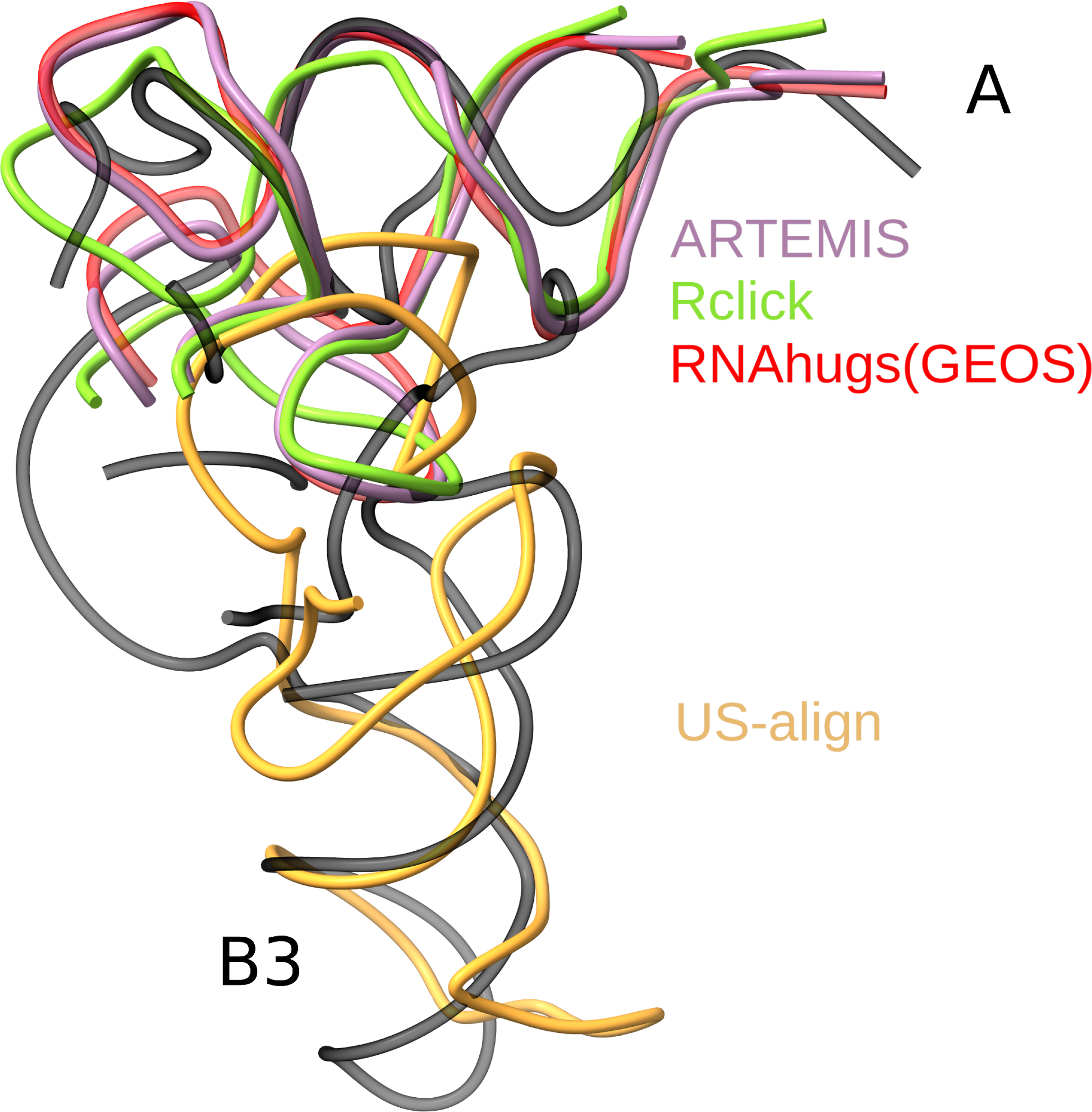
Topology-independent superposition of a tRNA (PDB entry 1IVS, chain C) and a BMV tRNA-like structure (PDB entry 7SAM, chain A, in black) conducted by ARTEMIS (in purple), US-align (in orange), Rclick (in green), and RNAhugs(GEOS) (in red). Unaligned structure fragments are hidden for clarity. Labels A and B3 denote the tRNA-mimicking fragments. The incorrect superposition by RNAhugs(GENS) is not shown.

### Analysis of identified helical packing motif

Among the 17 backbone-permuted structure pairs not involving the bacterial Y RNA, we observed six pairs to be noisy artifacts identified between two large RNAs, four pairs of structures trivially superimposed via a single helical fragment, and seven pairs featuring the well-known minor-groove/minor-groove helical packing motif [32], often formed with widespread A-minor interactions [33] and GNRA-tetraloop/receptor motifs [34] (Supplementary Table S4).

To analyze backbone variations of the observed motif we manually collected representative RNA 3D structures of 15 non-coding RNA families from Rfam [35] and one synthetic RNA, all featuring the helical packing motif (Table 2, Figure 7A). This set included various ribozymes and riboswitches with no obvious evolutionary relations, illustrating the motif’s intrinsic presence in RNA structures. Remarkably, the helices packed similarly within completely different backbone topologies (Figure 7B, C). Benchmarks of the topology-independent superposition tools based on the selected set of 16 RNAs confirmed the superior performance of ARTEMIS (Figure 7D, Supplementary Figure S2, Supplementary Table S6).

**Figure 7.**
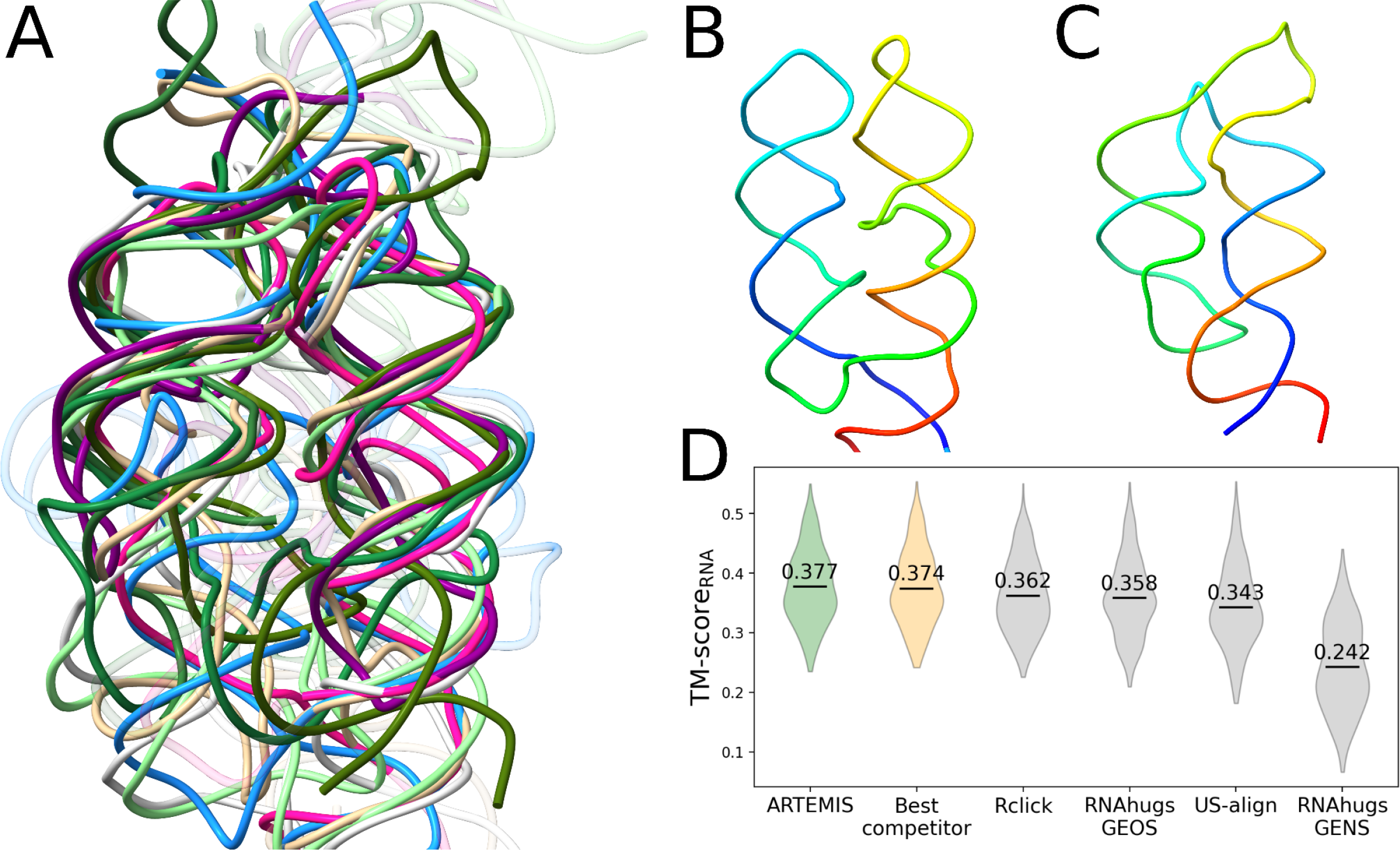
Minor-groove/minor-groove helical packing motif. (A) Topology-independent superposition constructed with ARTEMIS. (B) Archaeal SRP RNA (PDB entry 3NDB, chain M) and (C) THF riboswitch (PDB entry 6Q57, chain A) exhibiting the motif in distinct backbone topology contexts. The chains are rainbow-colored, beginning with blue at the 5′-end and ending with red at the 3′-end. (D) Comparisons of topology-independent superpositions built by different tools.

**Table 2.**
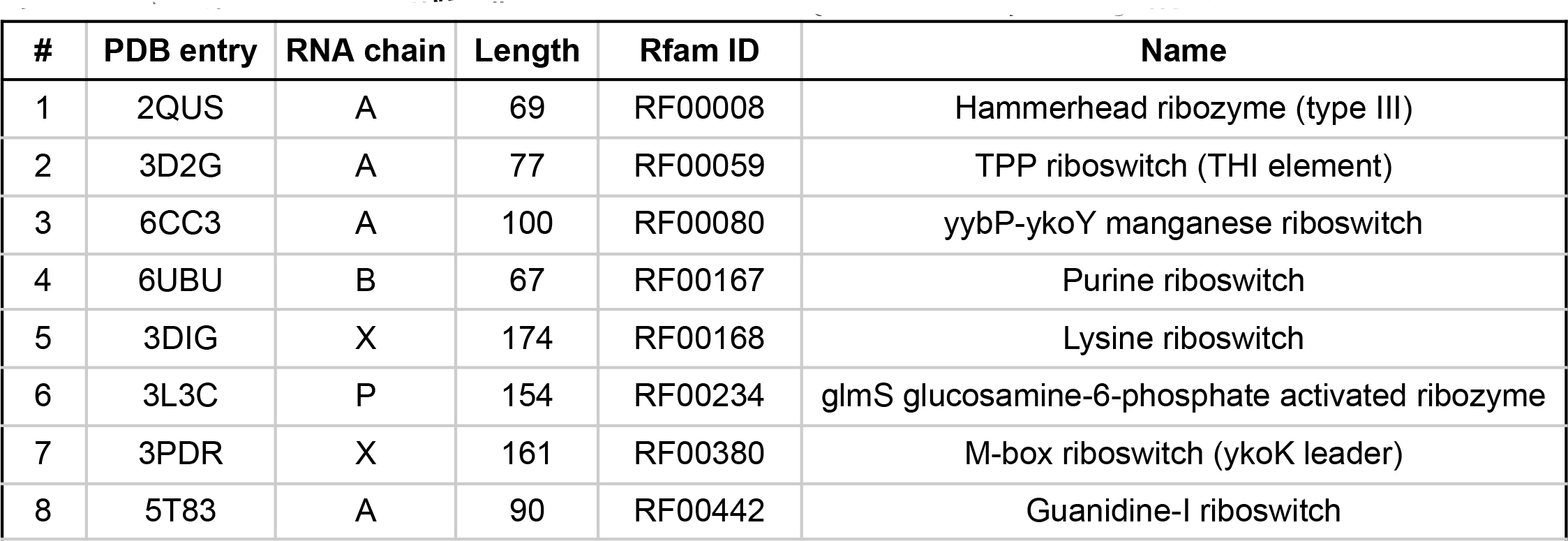
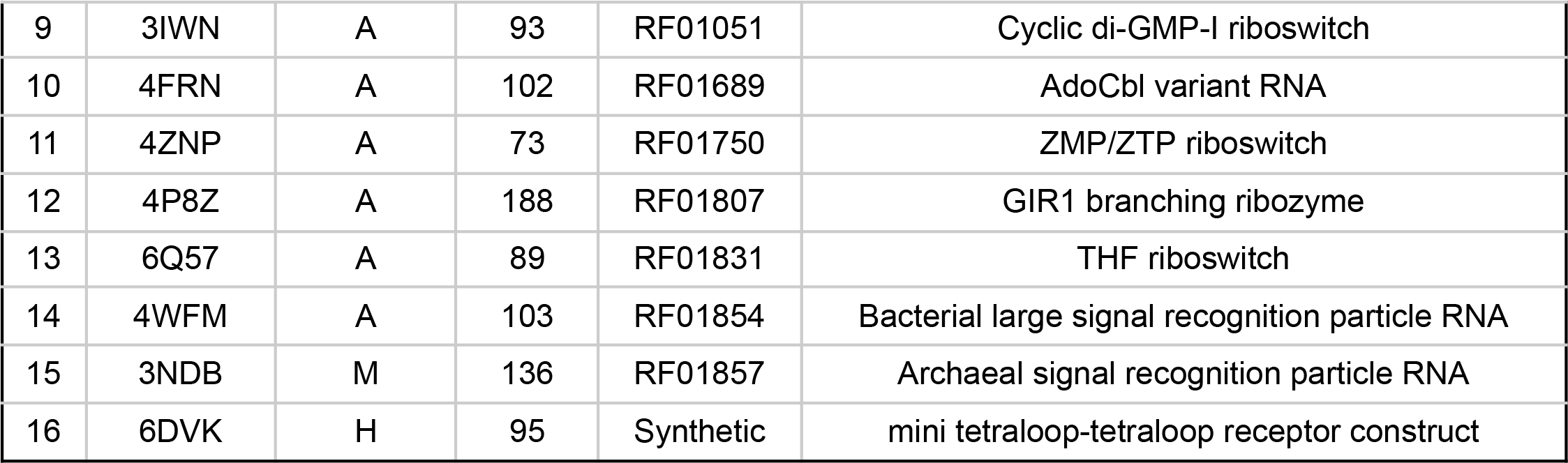
Representative RNA 3D structures featuring the helical packing motif.

## DISCUSSION

In this work, we introduced ARTEMIS, a new method for RNA 3D structure superposition. Our benchmarks demonstrate that ARTEMIS significantly outperforms state-of-the-art methods in both sequentially-ordered and topology-independent (potentially sequence-permuted) superpositions.

We utilized ARTEMIS to delve into the realm of backbone-permuted RNA structural similarities. The majority of such instances among the RNA 3D structures reported so far were tRNAs and tRNA-mimicking Y RNAs, alignable to each other with a circular permutation (Figure 5). Additionally, ARTEMIS successfully aligned tRNAs with viral tRNA-mimic structures, where the superpositions represent non-circular permutations (Table 1).

Through analysis of the RNA 3D structures with sequence permutations using ARTEMIS, we identified the minor-groove/minor-groove helical packing motif and showed that it is characteristic of multiple unrelated RNA molecules and can be formed within various backbone topology contexts (Figure 7). Based on our experience in RNA-Puzzles [36] and CASP15 [37, 38], such long-range interactions involving not just the individual residues but entire secondary structure elements are not yet properly captured by any of the RNA 3D modeling programs. ARTEM and ARTEMIS can capture recurring interactions in RNA 3D structures involving multiple local structural motifs, which may hold a key to understanding the folding of large RNA molecules that rely on long-range interactions.

Unlike other tools, ARTEMIS offers the capability for users to specify parts of the input structures to be used for superposition and separately designate which parts of the query coordinate file will be saved as the superimposed structure. This feature eliminates the need for manual editing of input coordinate files and proves useful e.g., for superimposing macromolecular complexes, including RNA-protein complexes, based on specified RNA chains, fragments, or even particular residues. ARTEMIS is also suitable for the superposition of DNA molecules, including G-quadruplexes of different topological arrangements, although for G-quadruplexes, Rclick performs comparably to ARTEMIS (data not shown). Furthermore, ARTEMIS can recognize all types of modified residues (e.g., according to [39]) if the user adds their atomic representations to the config files (*artemis_1.json* and *artemis_2.json* files, see https://github.com/david-bogdan-r/ARTEMIS/tree/main/src/resrepr).

We anticipate that ARTEMIS will find utility in various studies related to 3D structures of RNA, DNA, and nucleic acid-containing complexes. It will prove particularly valuable for the comparative analysis of RNA 3D folds and motifs featuring sequence permutations. Our plans include extending ARTEM/ARTEMIS to accommodate protein structures and ligands, reporting sub-optimal superpositions, performing multiple structure/sequence alignments, and flexible 3D structure superpositions.

## DATA AVAILABILITY

An implementation of the ARTEMIS tool and benchmarking data are available at https://github.com/david-bogdan-r/ARTEMIS (DOI: <u>10.5281/zenodo.10932928</u>).

## ACKNOWLEDGMENTS

The authors thank Dominik Sordyl for fruitful discussions and invaluable feedback.

## SUPPLEMENTARY MATERIALS

**Supplementary Figure S1.**
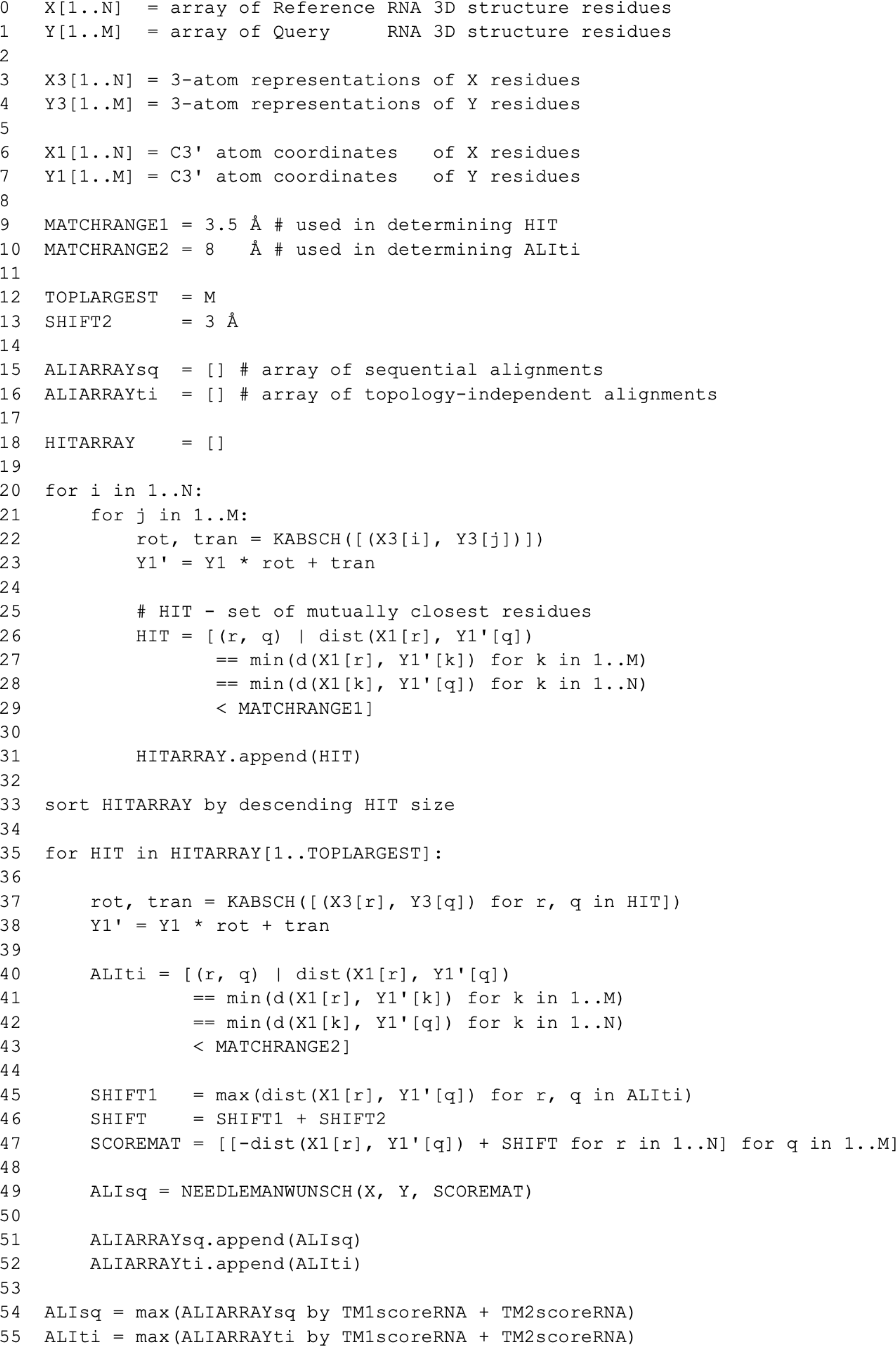
ARTEMIS algorithm pseudocode.

**Supplementary Figure S2.**
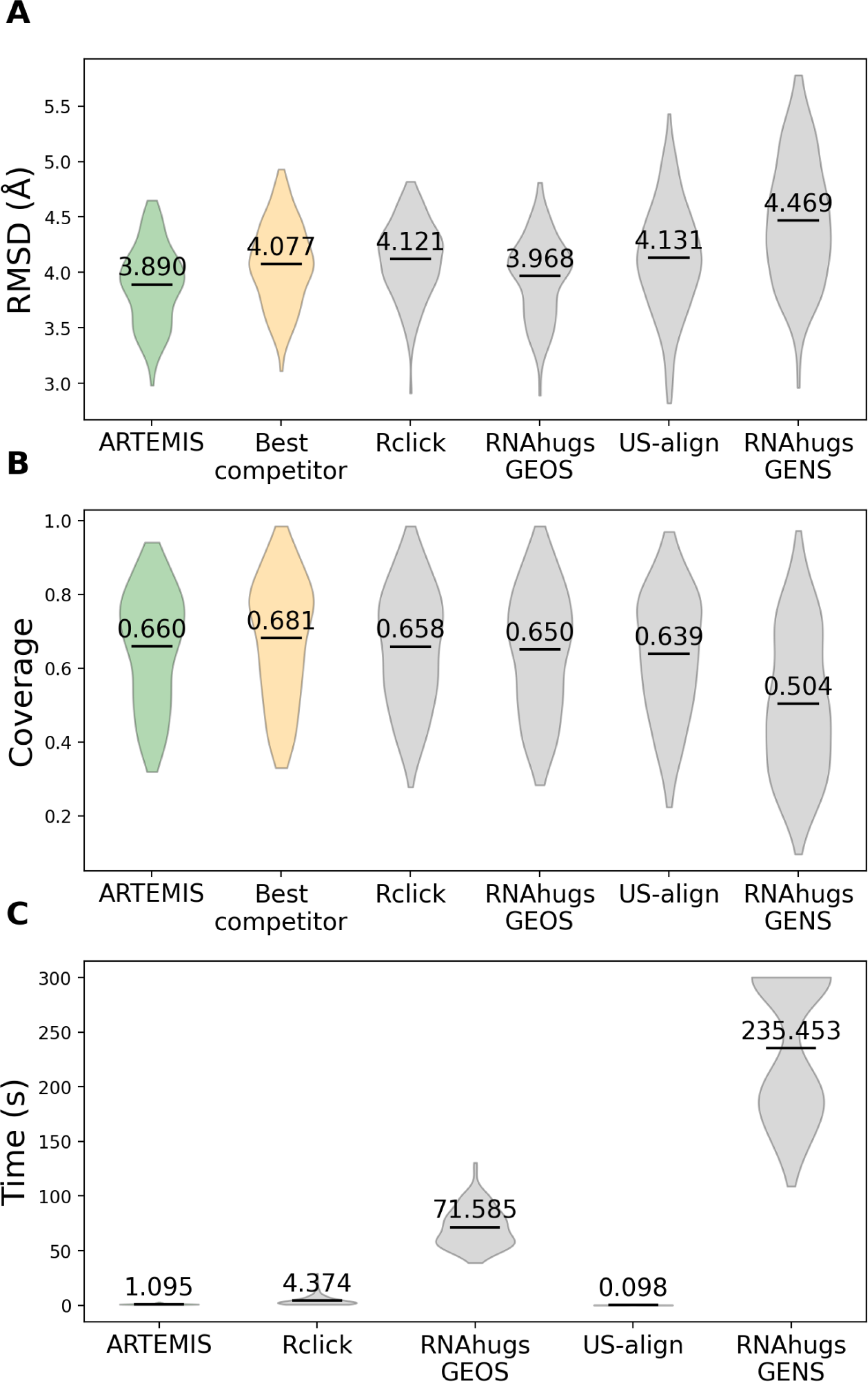
Topology-independent superposition comparisons for RNAs featuring the helical packing motif. Performance was measured by (A) RMSD, (B) coverage, and (C) execution time. ARTEMIS demonstrates superior performance compared to existing tools, as evidenced by lower RMSD values and higher coverage values.

**Supplementary Table S1.** Technical details of running the standalone versions of superposition tools.

| Tool | Description |  |
| --- | --- | --- |
| **Rclick** | Downloaded from | https://mspc.bii.a-star.edu.sg/minhn/download.html |
|  | Installation details | Changed the typeAtom parameter from CA to C3' in the Parameters.inp file |
|  | Running command | ./click inputPDB1 inputPDB2 |
| **RNAhugs** | Downloaded from | https://github.com/RNAPolis/rnahugs |
|  | Installation details | - |
|  | Command for GEOS mode | java -jar target/rna-hugs-1.0-jar-with-dependencies.jar --reference inputPDB1 --model inputPDB2 --method geometric |
|  | Command for GENS mode | java -jar target/rna-hugs-1.0-jar-with-dependencies.jar --reference inputPDB1 --model inputPDB2 --method genetic |
| **US-align** | Downloaded from | https://zhanggroup.org/US-align/bin/module/USAlign.cpp |
|  | Installation details | - |
|  | Command for topology-independent mode | ./USAlign inputPDB1 inputPDB2 -mm 5 |
|  | Command for topology-independent mode without re-superimposing input structures | ./USAlign superPDB1 superPDB2 -mm 5 -se |

**Supplementary Table S2.**
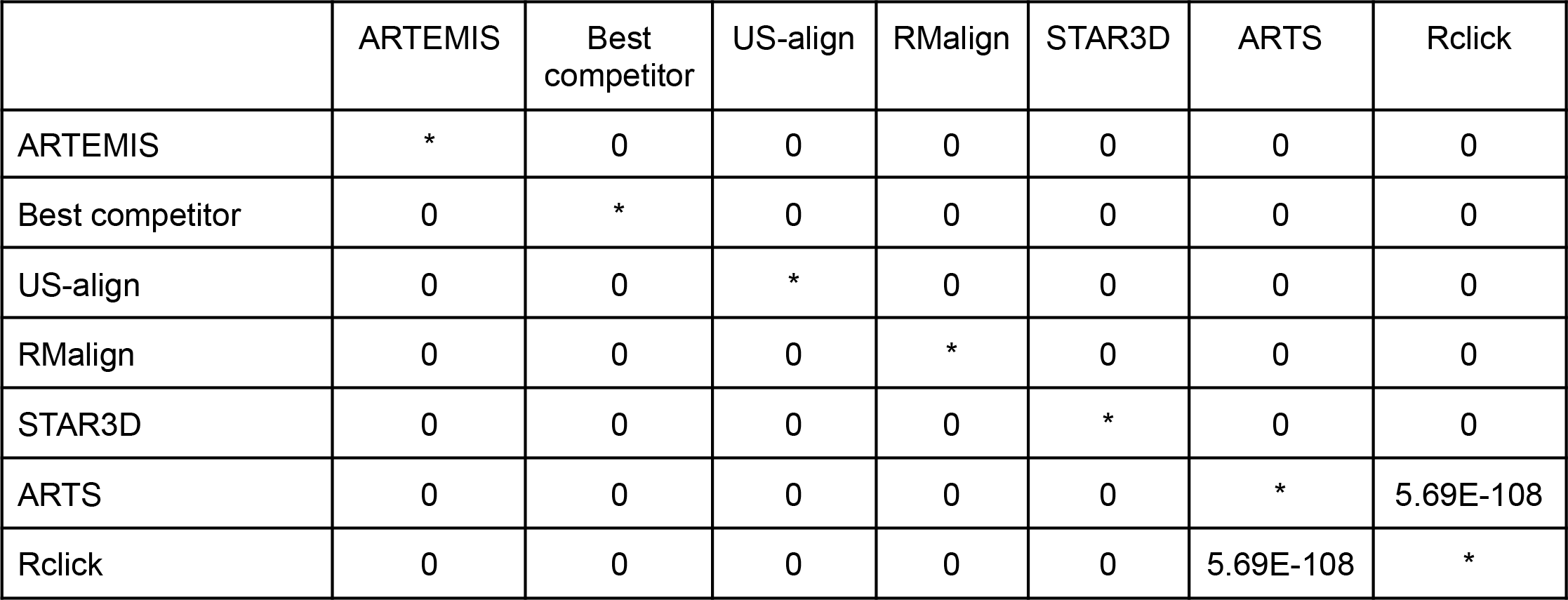
Student’s paired t-test p-values calculated for each pair of the superposition tools based on the sets of TM-score_RNA_ values, as measured on the dataset of 637 RNA chains in building sequentially-ordered alignments.

**Supplementary Table S3.** Student’s paired t-test p-values calculated for each pair of the superposition tools based on the sets of TM-score_RNA_ values, as measured on the dataset of 637 RNA chains in building topology-independent alignments.

|  | ARTEMIS | Best competitor | US-align | Rclick |
| --- | --- | --- | --- | --- |
| ARTEMIS | * | 0 | 0 | 0 |
| Best competitor | 0 | * | 0 | 0 |
| US-align | 0 | 0 | * | 0 |
| Rclick | 0 | 0 | 0 | * |

**Supplementary Table S4.** The list of 97 backbone-permuted structural similarities. Structure identifiers are built by concatenating PDB entries and chain identifiers. Seven pairs featuring the helical packing motif are underscored.

|  |  |  |  |  |
| --- | --- | --- | --- | --- |
| 6cu1A 6t4q6<br>6cu1A 6r87B<br>6cu1A 5o2rx<br>6cu1A 6t7t6<br>6cu1A 2derC<br>6cu1A 6r5q2<br>6cu1A 4wj4B<br>6cu1A 6q9a7<br>6cu1A 1qf6B<br>6cu1A 6tbvPTR1<br>6cu1A 6tb3n<br>6c4hA 6chrA<br>6cu1A 6ufmA<br>6cu1A 4v6uA0<br>6cu1A 5j8bx<br>6cu1A 2dr2B<br>6cu1A 3amtB<br>6cu1A 6i0yV<br>6cu1A 3j7a7<br>6cu1A 1j1uB | 5v93a 5z3gA<br>5hd12a 3j7yA<br>6cu1A 6t4q7<br>6cu1A 4v9iAY<br>6cu1A 4yyeC<br>6cu1A 6ip5zu<br>6cu1A 4v7IAY<br>5u4ja 2a64A<br>6cu1A 5wwrC<br>4v8dAB 6cu1A<br>6gawBA 4v4aAA<br>3q1qC 6cu1A<br>6cu1A 6cae1y<br>6cu1A 1il2C<br>6cu1A 5vppQV<br>6cu1A 6v3av<br>6cu1A 1gaxC<br>6cu1A 6az12<br>5g2xA 6c4hA<br>6cu1A 2csxC | 6cu1A 2zueB<br>6cu1A 3a2kC<br>6cu1A 6rfIU<br>6cu1A 3wc1P<br>5on2B 6cu1A<br>6cu1A 2du3D<br>6jxmB 6cu1A<br>6cu1A 3j46p<br>6cu1A 5e6mC<br>6cu1A 4rdxC<br>2zzmB 6cu1A<br>6cu1A 5b63D<br>6cu1A 6q977<br>4v8bAB 6cu1A<br>6cu1A 4v6uA1<br>6cu1A 2dlcY<br>6cu1A 1jgqD<br>6ah3T 6cu1A<br>1h3eB 6cu1A<br>6cu1A 6ek0S6 | 6cu1A 6uggA<br>6cu1A 4lckB<br>6cu1A 5x6bP<br>6cu1A 3wqyC<br>6cu1A 3ephE<br>6cu1A 5el63K<br>1wz2C 6cu1A<br>5ah5D 6cu1A<br>6cu1A 1h4sT<br>6cu1A 5lzsii<br>6cu1A 2fk6R<br>6cu1A 4v5IAY<br>6cu1A 2d6fF<br><u>3dilA 3pdrA</u><br>6cu1A 4v5gAY<br>6cu1A 5ud5D<br>6cu1A 2iy5T<br>6cu1A 1serT<br>6cu1A 3icqD<br>5lzdY 6cu1A | 6cu1A 2nreF<br>6cu1A 3akzE<br>6cu1A 4v8nAW<br><u>5tpyA 5t5aA</u><br>6cu1A 1u0bA<br>3am1B 6cu1A<br>6cu1A 1g59B<br>1s03A 1i6uC<br><u>1u9sA 3ndbM</u><br>3w1kF 6cu1A<br><u>6dmcA 3iwnA</u><br><u>3iwnA 5t83A</u><br><u>6dvkH 3iwnA</u><br><u>3iwnA 5u3gB</u><br>5m0hA 4pcjA<br>4jf2A 2j288<br>1m5oB 5ob3A |

**Supplementary Table S5.** Performance of the topology-independent superposition tools on tRNA-like structures in RMSD and coverage values.

| Tool |  |  |  | ARTEMIS |  | US-align |  | Rclick |  | RNAhugs<br>GEOS |  | RNAhugs<br>GENS |  |
| --- | --- | --- | --- | --- | --- | --- | --- | --- | --- | --- | --- | --- | --- |
| tRNA vs. | PDB id<br>Chain | Length | Permutation | RMSD | tRNA<br>Cov. | RMSD | tRNA<br>Cov. | RMSD | tRNA<br>Cov. | RMSD | tRNA<br>Cov. | RMSD | tRNA<br>Cov. |
|  | 1ivsC | 75 |  |  |  |  |  |  |  |  |  |  |  |
| tmRNA<br>fragment | 2czjB | 62 | no | 3.65 | 0.573 | 2.82 | 0.52 | 2.63 | 0.507 | 3.03 | 0.56 | 2.72 | 0.507 |
| tRNA-Sec | 3addC | 88 | no | 3.18 | 0.853 | 3.02 | 0.853 | 3.01 | 0.813 | 3.02 | 0.853 | 3.02 | 0.853 |
| TYMV<br>tRNA-like | 4p5jA | 83 | non-circular | 3.19 | 0.84 | 3.06 | 0.867 | 3.43 | 0.853 | 3.15 | 0.867 | 3.27 | 0.867 |
| Y RNA | 6cu1A | 79 | circular | 2.69 | 0.907 | 2.4 | 0.907 | 2.6 | 0.907 | 2.46 | 0.907 | 2.45 | 0.907 |
| BMV<br>tRNA-like | 7samA | 169 | non-circular | 4.15 | 0.853 | 4.26 | 0.707 | 4.18 | 0.88 | 4.05 | 0.76 | 3.55 | 0.587 |
| Mean |  |  |  | 3.372 | 0.805 | 3.112 | 0.771 | 3.17 | 0.792 | 3.142 | 0.789 | 3.002 | 0.744 |

**Supplementary Table S6.** Student’s paired t-test p-values calculated for each pair of the superposition tools based on the sets of TM-score_RNA_ values, as measured on the dataset of 16 RNA chains in building topology-independent alignments.

|  | ARTEMIS | Best competitor | Rclick | RNAhugs GEOS | US-align | RNAhugs GENS |
| --- | --- | --- | --- | --- | --- | --- |
| ARTEMIS | * | 0.00238 | 6.78E-29 | 4E-26 | 1.94E-30 | 8.42E-65 |
| Best competitor | 0.00238 | * | 1.62E-25 | 8.59E-25 | 6.16E-31 | 2.28E-61 |
| Rclick | 6.78E-29 | 1.62E-25 | * | 0.0495 | 4.54E-12 | 2.12E-54 |
| RNAhugs GEOS | 4E-26 | 8.59E-25 | 0.0495 | * | 2.09E-07 | 5.61E-52 |
| US-align | 1.94E-30 | 6.16E-31 | 4.54E-12 | 2.09E-07 | * | 3.72E-43 |
| RNAhugs GENS | 8.42E-65 | 2.28E-61 | 2.12E-54 | 5.61E-52 | 3.72E-43 | * |

